# Mucosal gene expression in response to SARS-CoV-2 is associated with early viral load

**DOI:** 10.1101/2022.08.23.504908

**Authors:** Seesandra V. Rajagopala, Britton A. Strickland, Suman B. Pakala, Kyle S. Kimura, Meghan H. Shilts, Christian Rosas-Salazar, Hunter M. Brown, Michael H. Freeman, Bronson C. Wessinger, Veerain Gupta, Elizabeth Phillips, Simon A. Mallal, Justin H. Turner, Suman R. Das

**Affiliations:** Department of Medicine, Vanderbilt University Medical Center, Nashville, TN; Pathology Microbiology and Immunology, Vanderbilt University Medical Center, Nashville, Tennessee, USA; Department of Otolaryngology, Vanderbilt University Medical Center, Nashville, TN; Department of Pediatrics, Vanderbilt University Medical Center, Nashville, TN

**Keywords:** Metatranscriptomic, RNAseq, mucosal immune response, gene expression, nasal swab, coronavirus, COVID-19, SARS-CoV-2

## Abstract

Little is known about the relationships between symptomatic early-time SARS-CoV-2 viral load and upper airway mucosal gene expression and immune response. To examine the association of symptomatic SARS-CoV-2 early viral load with upper airway mucosal gene expression, we profiled the host mucosal transcriptome from nasopharyngeal swab samples from 68 adults with symptomatic, mild-to-moderate COVID-19. We measured SARS-CoV-2 viral load using qRT-PCR. We then examined the association of SARS-CoV-2 viral load with upper airway mucosal immune response. We detected SARS-CoV-2 in all samples and recovered >80% of the genome from 85% of the samples from symptomatic COVID-19 adults. The respiratory virome was dominated by SARS-CoV-2, with limited co-detection of common respiratory viruses i.e., only the human Rhinovirus (HRV) being identified in 6% of the samples. We observed a significant positive correlation between SARS-CoV-2 viral load and interferon signaling (OAS2, OAS3, IFIT1, UPS18, ISG15, ISG20, IFITM1, and OASL), chemokine signaling (CXCL10 and CXCL11), and adaptive immune system (IFITM1, CD300E, and SIGLEC1) genes in symptomatic, mild-to-moderate COVID-19 adults, when adjusted for age, sex and race. Interestingly, the expression levels of most of these genes plateaued at a CT value of ~25. Overall, our data shows that early nasal mucosal immune response to SARS-CoV-2 infection is viral load dependent, which potentially could modify COVID-19 outcomes.

**AUTHOR SUMMARY:** Several prior studies have shown that SARS-CoV-2 viral load can predict the likelihood of disease spread and severity. A higher detectable SARS-CoV-2 plasma viral load was associated with worse respiratory disease severity. However, the relationship between SARS-CoV-2 viral load and airway mucosal gene expression and immune response remains elusive. We profiled the nasal mucosal transcriptome from nasal samples collected from adults infected with SARS-CoV-2 during Spring 2020 with mild-to-moderate symptoms using a comprehensive metatranscriptomics method. We observed a positive correlation between SARS-CoV-2 viral load with interferon signaling, chemokine signaling, and adaptive immune system in adults with COVID-19. Our data suggest that early nasal mucosal immune response to SARS-CoV-2 infection was viral load-dependent and may modify COVID-19 outcomes.

## INTRODUCTION

Severe Acute Respiratory Syndrome Coronavirus-2 (SARS-CoV-2) infection causes Coronavirus Disease-19 (COVID-19) and is responsible for the 21^st^ century’s most significant pandemic [1]. While the majority of SARS-CoV-2 infections are either mild or asymptomatic, in approximately 2%-10% cases, it can lead to life threatening pneumonia and multiple organ failure [2], resulting in more than 5.5 million official COVID-19 deaths worldwide[3]. It has been hypothesized that a dysregulated innate immune activation response induces a cytokine storm that promotes respiratory failure and may lead to acute respiratory distress syndrome (ARDS) [4, 5], which has been the main reason for hospital admission and mortality in COVID-19 patients [5]. Several studies suggested that SARS-CoV-2 viral load can predict the likelihood of disease spread and severity [6-8]. A higher detectable SARS-CoV-2 plasma viral load was associated with worse respiratory disease severity [8]. Recently, we have identified associations between SARS-CoV-2 viral titer and the nasopharyngeal microbiome in adults [9]. However, little is known about the relationship between SARS-CoV-2 viral load and airway mucosal immune response. To fill this gap in knowledge, we profiled the host mucosal transcriptome from nasopharyngeal samples collected from outpatient adults infected with SARS-CoV-2 during Spring 2020 with mild-to-moderate symptoms utilizing a comprehensive metatranscriptomic method we recently developed [10]. We examined the association of SARS-CoV-2 viral load with the mucosal immune response.

## RESULTS

### Patients Characteristics and SARS-CoV-2 viral load measurement

Sixty-eight adults with confirmed, symptomatic, mild-to-moderate COVID-19 [based on criteria from the World Health Organization [11]] were enrolled as part of a clinical trial [12]. All samples patient enrollment and sample collection were done in the early phase of the pandemic (spring of 2020). As described in the methods, all research samples were collected early in the infection (within 24 hrs. of clinical diagnosis). The patients baseline characteristics are shown in **Table 1**. The median (interquartile range [IQR]) age was 36 (27-57) years. None of the participants had used antibiotics in the prior two weeks. Quantitative reverse transcription PCR (RT-qPCR) was used for quantification of SARS-CoV-2 viral load. Based on the cycle threshold (CT) value, we partitioned the subjects into tertiles: lower tertile (CT 14.5-21.5, n=23, hereinafter referred to as “high viral load group”), median tertile (CT 21.6-25.5, n=22, hereinafter referred to as “medium viral load group”), and upper tertile (CT 25.6 – 36, n=23, hereinafter referred to as “low viral load group”), respectively. There were no significant differences in baseline characteristics and underlining comorbidities between the three groups (**Table 1**).

**Table 1.** Demographic characteristics by viral load tertiles.

|  | High viral load<br>N=23 | Medium viral load<br>N=22 | Low viral load<br>N=23 | Combined<br>N=68 | Test Statistic |
| --- | --- | --- | --- | --- | --- |
| Age | 30 46 59<br>44 ± 16 | 24 30 45<br>35 ± 15 | 28 36 58<br>41 ± 16 | 27 36 57<br>40 ± 16 | $F_{2,65}=2.1$, $P=0.13^1$ |
| Sex : Male | 0.43 $\frac{10}{23}$ | 0.45 $\frac{10}{22}$ | 0.61 $\frac{14}{23}$ | 0.50 $\frac{34}{68}$ | $\chi^2_2=1.7$, $P=0.44^2$ |
| Race: African American | 0.04 $\frac{1}{23}$ | 0.05 $\frac{1}{22}$ | 0.22 $\frac{5}{23}$ | 0.10 $\frac{7}{68}$ | $\chi^2_6=11$, $P=0.082^2$ |
| Hispanic | 0.00 $\frac{0}{23}$ | 0.09 $\frac{2}{22}$ | 0.04 $\frac{1}{23}$ | 0.04 $\frac{3}{68}$ | |
| Unknown | 0.30 $\frac{7}{23}$ | 0.09 $\frac{2}{22}$ | 0.09 $\frac{2}{23}$ | 0.16 $\frac{11}{68}$ | |
| White | 0.65 $\frac{15}{23}$ | 0.77 $\frac{17}{22}$ | 0.65 $\frac{15}{23}$ | 0.69 $\frac{47}{68}$ | |
| Diabetes | 0.09 $\frac{2}{23}$ | 0.09 $\frac{2}{22}$ | 0.04 $\frac{1}{23}$ | 0.07 $\frac{5}{68}$ | $\chi^2_2=0.46$, $P=0.79^2$ |
| Hypertension | 0.26 $\frac{6}{23}$ | 0.14 $\frac{3}{22}$ | 0.17 $\frac{4}{23}$ | 0.19 $\frac{13}{68}$ | $\chi^2_2=1.2$, $P=0.55^2$ |
| Heart disease | 0.04 $\frac{1}{23}$ | 0.00 $\frac{0}{22}$ | 0.09 $\frac{2}{23}$ | 0.04 $\frac{3}{68}$ | $\chi^2_2=2$, $P=0.36^2$ |
| Pulmonary disease | 0.13 $\frac{3}{23}$ | 0.18 $\frac{4}{22}$ | 0.09 $\frac{2}{23}$ | 0.13 $\frac{9}{68}$ | $\chi^2_2=0.88$, $P=0.64^2$ |
| obese | 0.31 $\frac{5}{16}$ | 0.26 $\frac{5}{19}$ | 0.40 $\frac{8}{20}$ | 0.33 $\frac{18}{55}$ | $\chi^2_2=0.85$, $P=0.65^2$ |
| a b c represent the lower quartile a, the median b, and the upper quartile c for continuous variables. $\bar{x} \pm s$ represents $\bar{X} \pm 1$ SD. N is the number of non-missing values. Tests used: <sup>1</sup>Kruskal-Wallis test; <sup>2</sup>Pearson test. | | | | | |

### Metatranscriptome captured the nasal virome

Total RNA from the nasopharyngeal swab samples was extracted and further processed as previously described [10]. After quality-based trimming and removal of human and bacterial rRNA reads, we had an average of 98,120,791 (740,58,180 – median) human reads and an average of 2,263,290 (848,478 – median) microbiome reads. The microbiome reads bin contains viral, bacterial, fungal, and unclassified reads (for details see methods). Following quality control and initial data processing steps including taxonomic classification of reads, the reads classified as viruses were used to profile the respiratory virome and the reads that mapped to human transcripts were used to analyze the host response to SARS-CoV-2.

With a stringent cut-off of >80% genome coverage, we identified SARS-CoV-2 in 58 (85.29%) out of 68 samples. All SARS-CoV-2 genomes were identified were placed to the B.1 lineage in the Nextclade tree [13]. In addition to SARS-CoV-2, we identified HRV in four samples, and bacteriophage/prophage RNA transcripts were also recovered from many of the samples, albeit with lower genome coverage (**Figure 1a**). Surprisingly, common respiratory viruses were not co-detected in these samples, except HRV was co-detected in 6% of the samples. This may be due to implementation of public health measures, like social distancing and masking practices because of the COVID-19 early stage of the pandemic.

**Figure 1:**
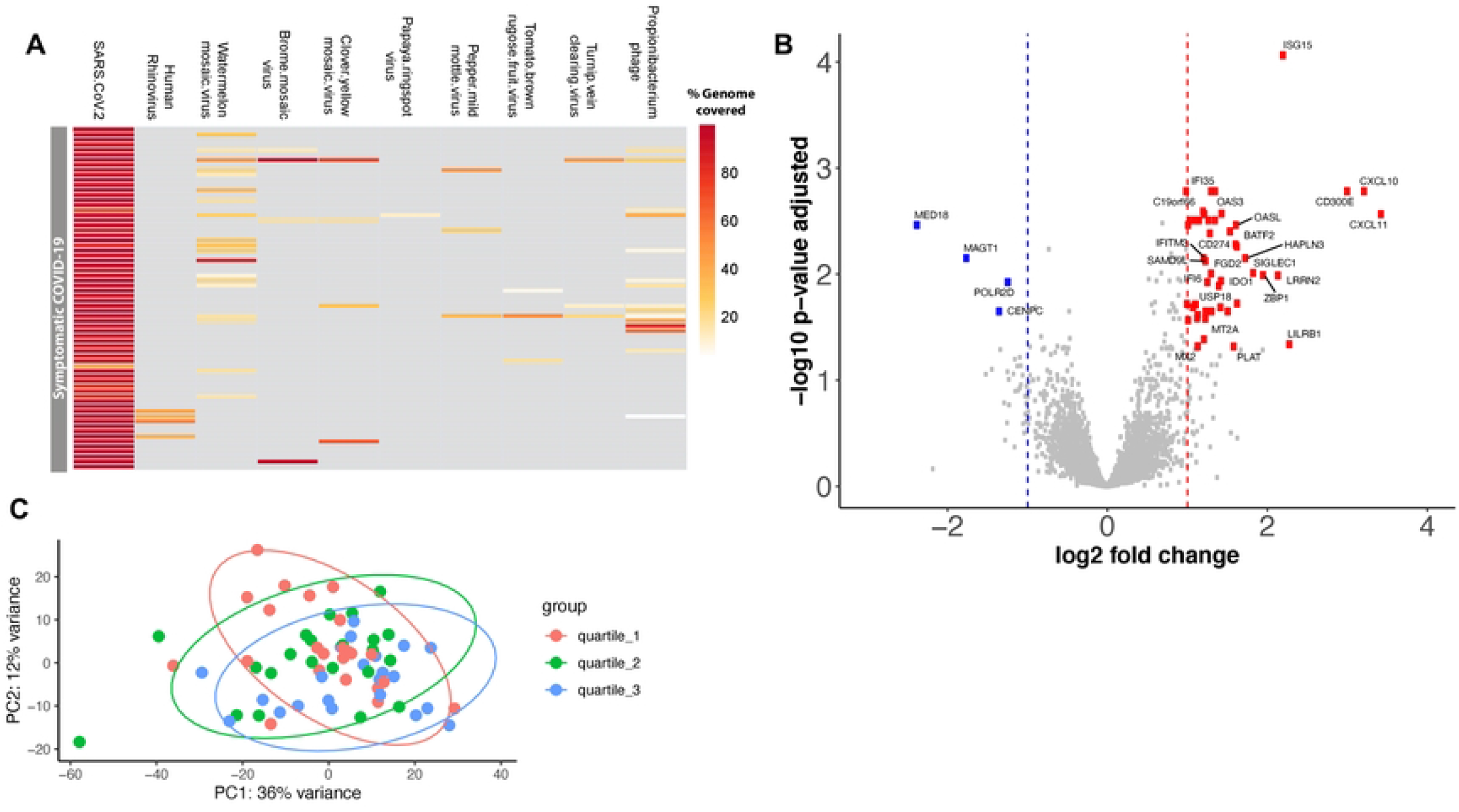
Differentially expressed genes between high and low viral load SARS-COV-2 infected adults. (a) Heatmap showing the virome profile. Each row represents a sample and each column represents the percentage of a virus genome recovered. (b) Volcano plot showing of log2 fold change and adjusted p-value obtained from DESeq2 analyses. Differential expression analysis was conducted using DESeq2 models, including age, sex, and ethnicity as covariates. The red circles indicate significantly up-regulated genes, and blue circles indicate significantly down-regulated genes in symptomatic SARS-COV-2 adults with high viral load compared to the low viral load group. Only the top 50 most significantly different genes are labeled. (c) Principal component analysis of the normalized gene-level read counts. Each dot represents a sample, and the samples are colored based on the CT value; we partitioned the subjects into tertiles: lower tertile (high viral load group) shown in red, median tertile (medium viral load group) shown in green, and upper tertile (low viral load group) shown in blue color.

### Differential host nasal mucosal gene expression based on SARS-CoV-2 viral load

Differential expression analysis between high and low viral load groups revealed 48 genes were significantly upregulated and 4 genes were significantly downregulated (based on a threshold of log2 fold change > |1| and adjusted p < 0.05) in the high viral load subjects (**Figure 1b-c and Supplementary Table 1**). Most of the upregulated genes in the high viral load group are involved in the immune response during viral infection, specifically genes involved in interferon alpha/beta signaling (IFITM3, IFITM1, RSAD2, MX2, IFI6, ISG15, IFI35, IFIT1, USP18, OASL, BST2, ISG20, OAS1, OAS2, OAS3, IRF7, XAF1) (**Figure 2a-b**) and immunoregulatory interactions between a lymphoid and a non-lymphoid cell (IFITM1, CD300E, LILRB1, SIGLEC1) (**Figure 2b**). Only a few genes were downregulated in the high viral load group, including MAGT1 (magnesium transporter 1), MED18 (mediator complex subunit 18), POLR2D (RNA polymerase II subunit D), and CENPC (centromere protein C) (**Figure 1**). There was no significant difference in the gene expression profiles between the high and medium viral load groups and medium and low viral load groups.

**Figure 2.**
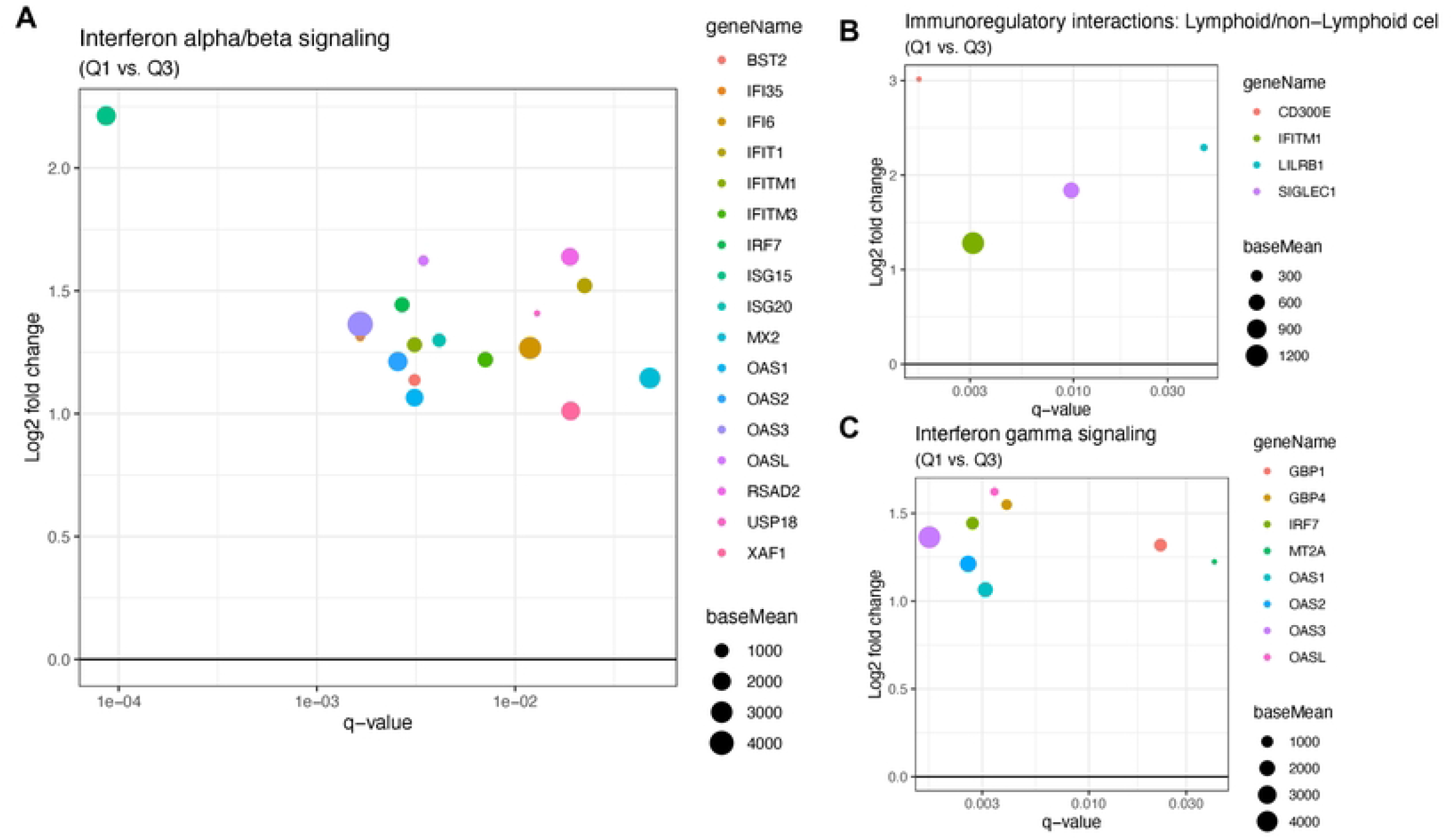
(a) Interferon alpha/beta signaling pathway genes that are significantly down- or up-regulated regulated in symptomatic SARS-COV-2 adults with high viral load compared to the low viral load group. On the x-axis is displayed the q-value for the up- or down-regulated genes with q-values < 0.05. On the y-axis is displayed the log2 fold change for those genes. The size of the dots represents “base mean” which is the mean of normalized counts of all samples. b) Similar to **(a)**, a plot showing the genes involved in Immunoregulatory interactions between a Lymphoid and a non-Lymphoid cell that are up-regulated in the high viral load group.

We examined the Spearman correlations between SARS-CoV-2 viral load and the mucosal gene expression levels. The genes that are involved in mucosal inflammation were inversely correlated to viral loads in terms of the CT values (**Figure 3**); a lower CT value corresponds to a high viral load, indicating a higher level of infectiousness. Strikingly, there was a moderate negative correlation (ρ between -0.5 to -0.6; q-value < 0.05) between interferon signaling (OAS2, OAS3, IFIT1, UPS18, ISG20, IFITM1, ISG15, and OASL), chemokine receptor signaling (CXCL 10 and CXCL11), and adaptive immune system (IFITM1, CD300E, and SIGLEC1) genes with SARS-CoV-2 CT values in symptomatic, mild-to-moderate COVID-19 adults (**Figure 3**). The expression levels of most of these genes were decreased with a lower viral load and seemed to reach the plateau phase at a CT value of ~25 (**Figure 4**). The expression of the POLR2D gene, which encodes the fourth largest subunit of RNA polymerase II that is responsible for synthesizing messenger RNA in eukaryotes, showed weak positive correlation (*r* 0.28, q-value: 0.02) with SARS-CoV-2 viral load (**Figure 3**).

**Figure 3:**
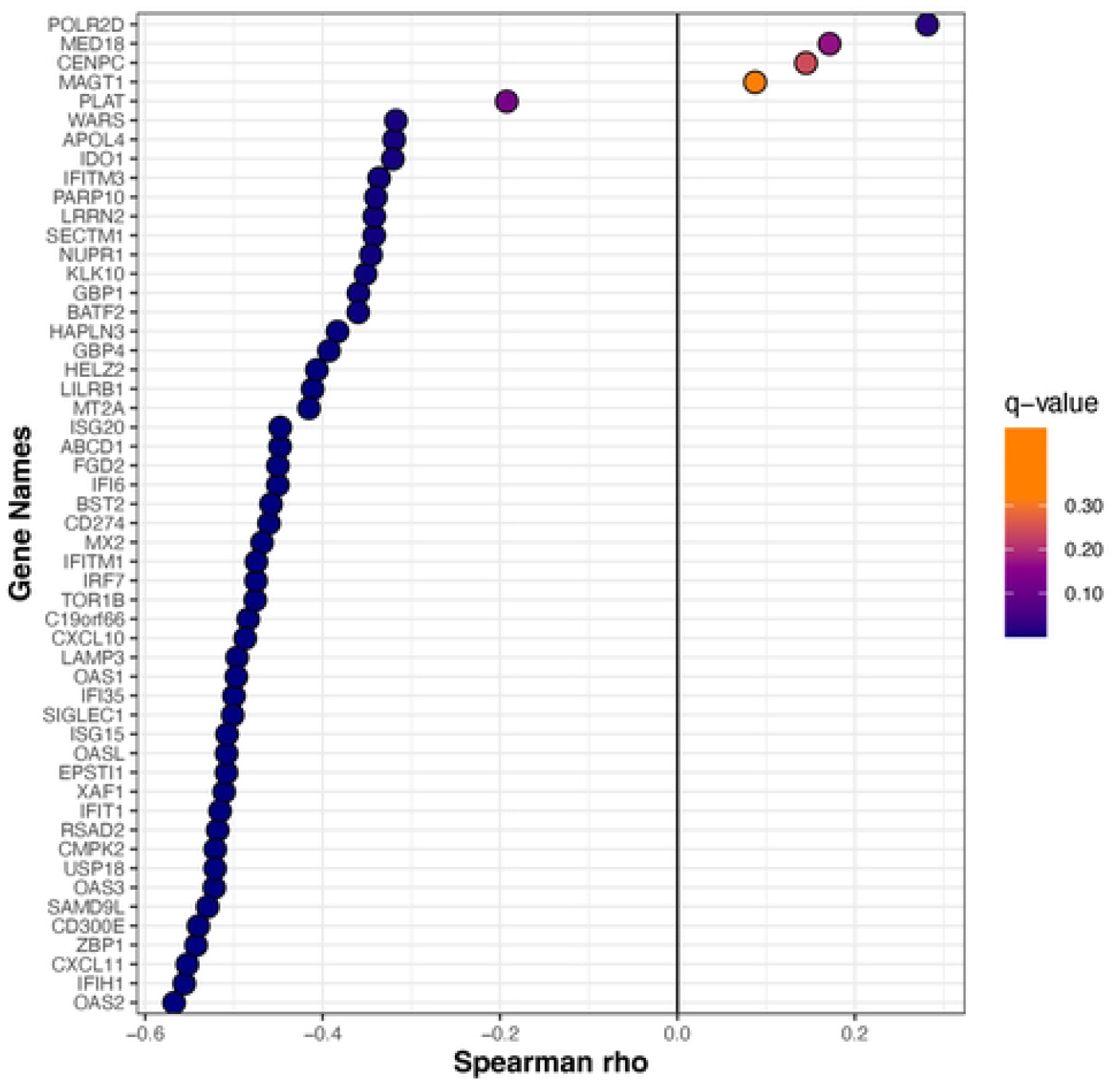
The Spearman correlation between CT value and mucosal gene expression. Spearman’s correlation coefficient was calculated for normalized gene-level expression counts and CT value (viral load). The Benjamini-Hochberg method was used for multiple comparisons and p-value adjustments. The gene expression is moderately correlated with viral load. As the *adjusted-p* < 0.05, the correlation is statistically significant.

**Figure 4:**
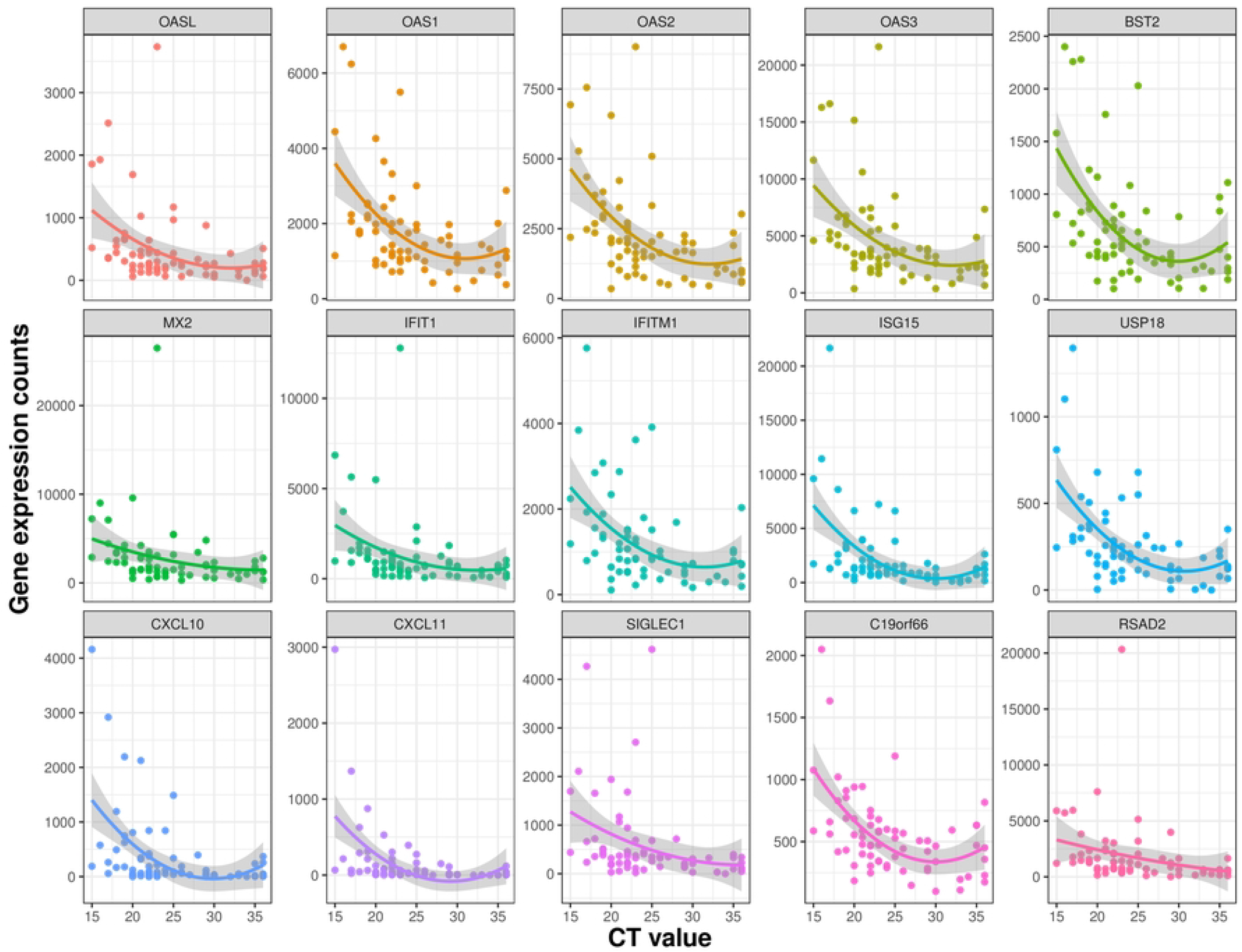
Scatter plot showing a correlation between CT value and mucosal gene expression. Normalized expression counts for selected genes of all samples were plotted with the CT values. The expression levels of most of these genes decreased with a lower viral load (i.e., with increased CT values) and reach the plateau phase at a CT value of ~25.

## DISCUSSION

High SARS-CoV-2 viral loads during early infection have been associated with severe disease outcomes [14]. We did not find an association between disease severity and viral load in our cohort as all of the patients had mild-to-moderate symptoms. A multitude of studies have focused on the change in immune response and gene expression in human samples, including lung tissues, [15] tracheal aspirates, [16] nasal and nasopharyngeal swabs,[17-20], BAL fluid,[20] and blood[19, 21-24]. Studies using animal models and cell lines have shown that SARS-CoV-2 infection is associated with low type I and type III interferon and elevated IL6 and chemokines that resulted in reduced innate antiviral defense and increased inflammation [25]. A metatranscriptomic analysis of human BAL fluid from SARS-CoV-2-infected patients show robust overexpression of interferon stimulated genes (ISGs) and other inflammatory genes [20]. However, to our knowledge, none of the studies were focused on understanding the role of early viral load on mucosal immune response.

Similar to published data we found the antiviral protein ISG15 was upregulated upon SARS-CoV-2 infection [26], as it can activate downstream pattern recognition receptor MDA5 to establish an antiviral state. However, this pathway has been shown to be directly antagonized by SARS-CoV-2 to evade host innate immunity via overexpression of PLpro, a gene with redundant functionality of ISG-deconjugating enzyme USP18 (a negative regulator of type I and III interferons) [27], as several studies [27, 28] show upregulation of USP18 upon SARS-CoV-2 infection. Here we show USP18 was upregulated in the nose in a viral load dependent manner, which for the first time confirms *in vitro* findings in clinical samples (**Figure 4**). Our findings in the human clinical samples support the hypothesis from Vere et al. that targeting USP18 has strong therapeutic potential for COVID-19 [29].

Similar to our findings, several studies have shown that the ISGs, OAS1, OAS2, OAS3, IFITMs, and OASL are robustly activated upon SARS-CoV-2 infection [30, 31]. ISGs are critical in countering other interferon responsive viruses like SARS-CoV, and MERS-CoV [32]. Supporting this hypothesis, multiple genome wide association studies have shown that mutations in ISGs are associated with susceptibility to SARS-CoV-2 and severity of COVID-19. Similarly, we also found interferon alpha and beta signaling genes BST2, IFIT1, OASL, MX2 (known antivirals), IFI35, IFI6, IFITM1, IFIM3, IRF7, ISG15, ISG20, RSAD2, USP18, XAF1 (**Figure 4**) were upregulated in subjects with a higher viral load and many of these genes have been linked with COVID-19 or other viral infection restriction. IFIT genes, particularly IFIT1, work in concert with other ISGs to inhibit viral translation by binding viral mRNAs that mimic host mRNAs due to presence of 5’ cap-1 structures [33, 34], which SARS-CoV-2 contains [35]. IFIT proteins also increase the expression of CXCL10,[36] the 2^nd^ most upregulated gene in our analysis, which induces lymphocyte chemotaxis and may inhibit the replication of infected viruses. Only one study has identified IFIT1 during COVID-19 *in vitro* [37].

CXCL10 and CXCL11 are two T helper type 1 (Th1) chemokines that were both upregulated in our analysis, and their gene expression was corelated with viral load. Both chemokines are highly produced in bronchial and alveolar epithelial cells and have been implicated as predictors of COVID-19 severity *in vitro, in vivo* in animal model, and clinical data [38, 39]. Additionally, our findings of upregulation of RSAD2, OAS1-3, IFITM1, MX2, and CD300E [20, 40-42] agree with findings previously published for SARS-CoV-2 infection. We further show, the immune modulatory gene sialic acid-binding Ig-like lectin 1 (Siglec-1/CD169), was upregulated in subjects with high viral load. Lectins, especially Siglec-1/CD169, have been known to mediate the attachment of viruses to APCs [43]. Recent studies show blockage of Siglec-1 on monocyte-derived dendritic cells (MDDCs) decreased SARS-CoV-2 viral transfer or *trans*-infection to bystander target cells [43] and Siglec1 was associated with disease severity [44] and enhanced SARS-CoV-2 infection and influenced neutralizing antibodies [45].

Our study has many strengths, such as the inclusion of an overall young population with no underlining comorbidities (**Table 1**), no recent use of antibiotics or current use of intranasal medications, and samples were collected early in the infection, i.e., within the first 24 hours of SARS-CoV-2 diagnosis. Also, as the patient enrollment and sample collection were done in the early pandemic (spring 2020), none of our patients were vaccinated or potentially infected prior to this infection, which could confound immune response and gene expression [9, 12]. All the SARS-CoV-2 genotypes in this study were identified as B.1 lineage (Supplementary Table 2). Our metatranscriptomics method also captured the virome and shows limited co-infection of other respiratory viruses.

We should also acknowledge several limitations. First, our study was cross-sectional; thus, there is a possibility of reverse causation. Second, there is a possibility of residual confounding, as we lacked data on the participants’ atopic status or other unknown morbidities that could modify immune response. Third, though all samples were collected within 24 hours after the first diagnosis of COVID-19, it is still impossible to ascertain the days since infection as the pre-symptomatic period can vary considerably (3-15 days) between patients. Nasal viral titers drop dramatically after 10-15 days of infection [46]; thus, prior studies in hospitalized patients rather have shown blood viral titer to be better predictor of the disease severity [8]. Fifth, we did not have lower respiratory tract samples. However, the URT is the portal of entry and an active site of replication of SARS-CoV-2, as well as a common harboring site for potential pathogens, thus of critical importance in the pathogenesis of this respiratory virus.[47] Last, our results cannot be extended to adults with asymptomatic, severe, or critical COVID-19, as only those with symptomatic, mild-to-moderate COVID-19 were included in our study.[11] In spite of these limitations, our study highlights URT mucosal innate immune response correlates with SARS-CoV-2 viral load.

In summary, we determined early SARS-CoV-2 viral load was associated with URT gene expression during COVID-19 and potentially modifies both the innate and adaptive immune response to SARS-CoV-2 infection. Future studies with larger sample sizes, serial sample collection, and with patients who develop severe disease outcomes will be needed to examine how SARS-CoV-2 interacts with the mucosal gene expression and how these viral-host interactions can impact the clinical progression, severity, and recovery of COVID-19.

## MATERIALS AND METHODS

### Overview of the Study Design

For the current study, we included non-hospitalized patients aged ≥18 years who were diagnosed with severe acute respiratory syndrome coronavirus-2 (SARS-CoV-2) infection (confirmed by qualitative polymerase chain reaction [PCR]) at Vanderbilt University Medical Center or one of its affiliated centers in Nashville, Tennessee. These patients were enrolled as part of a clinical trial examining the effect of several types of nasal irrigations on upper respiratory tract (URT) symptoms and viral load during coronavirus disease 2019 (COVID-19). The detailed methods for this clinical trial have been previously reported.[12] Exclusion criteria for these patients included current use of nasal saline irrigations or other intranasal medications, inability to perform nasal irrigations or to collect URT samples in a separate house bathroom or away from household contacts, or need for hospitalization related to SARS-CoV-2 infection. Thus, only patients with mild or moderate COVID-19 (based on criteria from the World Health Organization[11]) were included in the clinical trial. Eligible patients were contacted and enrolled in the study within 24 hours of initial diagnosis. Following adequate training, all participants were asked to obtain a mid-turbinate swab on the day of enrollment (i.e., prior to any study intervention for those enrolled in the clinical trial) using a self-collection kit (FLOQSwabs, Copan Diagnostics Inc.). These enrollment samples were used for the current study. The collection of all samples included in the current study occurred between April and June of 2020. Each adult provided informed consent for their participation. The Institutional Review Board of Vanderbilt University approved this study.

### Severe Acute Respiratory Syndrome Coronavirus-2 Testing by Quantitative Reverse Transcription Polymerase Chain Reaction

To measure viral load in SARS-CoV-2-infected patients, we performed quantitative reverse transcription PCR (RT-qPCR) in the mid-turbinate swabs. Total RNA was extracted from the swabs using a phenol-chloroform-based method. The swabs were placed in Red 1.5 mL RINO^®^ screw-cap tubes (NextAdvance) pre-filled with RNase-free zirconium oxide beads and QIAzol Lysis Reagent (Qiagen) was added. Samples were then homogenized in a Bullet Blender 24 Gold (NextAdvance) for 3 minutes at maximum speed. Following homogenization, genomic DNA was eliminated with gDNA Eliminator columns (Qiagen) and RNA purified using the RNeasy Mini Plus Kit (Qiagen) following the manufacturer’s protocol. The RNA quality was measured using an Agilent 2100 Bioanalyzer (Agilent Technologies). The United States Centers for Disease Control and Prevention primers and probes designed for the detection of SARS-CoV-2 (2019-nCoV) were purchased from Integrated DNA Technologies (IDT)[48]. Both the SARS-CoV-2 nucleocapside gene region 1 (N1) and nucleocapside gene region 2 (N2) were targeted for the detection of SARS-CoV-2. RNase P was also examined as a measure of RNA quality and quantity. RT-qPCR was performed using SuperScript III One-Step RT-PCR System with Platinum Taq DNA Polymerase (Invitrogen) as per manufacturer’s instructions on a Bio-Rad CFX96 Touch Real-Time PCR Detection System (Bio-Rad). Plasmid controls for 2 SARS-CoV-2 nucleocapsid and RNase P were also ordered from IDT at a concentration of 66,666 copies/reaction. No template controls and an extraction negative were used as negative controls. Reactions were prepared using 12.5 μl SuperScript III Master Mix (ThermoFisher), 1 μl each 400nm forward and reverse primer, 1 μl 400nM FAM-labelled probe, 1 μl Platinum Taq polymerase, 3 μl of template RNA, and 7.25 μl PCR Certified Water (Teknova). RNA was reverse transcribed at 50ºC for 15 minutes, and PCR conditions were run at 95ºC denaturation step for 2 minutes, followed by 40 cycles of 95ºC for 15 seconds and 55ºC for 30 seconds. The cycle threshold (CT) values were captured and calculated by the CFX Maestro (Bio-Rad) software and used as a measure of viral load.

### RNA extraction, metatranscriptomic library preparation and sequencing

The nasal swab samples in the self-collection kit (FLOQSwabs, Copan Diagnostics Inc.) were vortexed for 2 minutes, then an aliquot of 250µL nasal swap sample was used for RNA extraction. The total RNA from these samples were extracted as describe previously [10]. In brief, 250µL aliquot from the nasal swab sample was homogenized in 600µL QIAzol (Qiagen) and 500µL of 2.0 mm zirconium oxide beads (Next Advance, Inc. Cat: ZROB05) using a Bullet Blender homogenizer (BB24-AU, Next Advance, Inc). While homogenizing the samples, temperature was maintained at or near 4°C by using the dry ice cooling system in the Bullet Blender. The homogenate was treated with 100µL of genomic DNA Eliminator solution (Qiagen) to remove the genomic DNA. Next, 180µL of chloroform was added to the samples for phase separation. The total RNA in the aqueous phase was then purified using RNeasy Mini spin columns as recommend by the Qiagen RNeasy protocol. RNA integrity and RNA quantification were assessed using an Agilent Bioanalyzer RNA 6000 Nano/Pico Chip (Agilent Technologies, Palo Alto, California). Eukaryotic ribosomal RNAs (rRNA) were depleted using the NEBNext rRNA Depletion Kit (Human/Mouse/Rat, Cat: E6310X). After rRNA depletion, the samples were checked by Agilent Bioanalyzer RNA 6000 Nano/Pico Chip to ensure depletion of the 18S and 28S ribosomal peaks. Next, Illumina sequencing libraries were made using the NEBNext Ultra II RNA Library Prep Kit (NEB #E7775). The quality of the libraries was assessed using an Agilent Bioanalyzer DNA High Sensitivity chip. The libraries were then sequenced on an Illumina NovaSeq6000 platform (S4 flow cells run) with 2×150 base pair reads, with a sequencing depth of ~40 million paired-end reads per sample.

### Quantification and statistical analysis

#### Preprocessing and quality control of NGS data

Adapter removal and quality-based trimming of the raw reads were performed using Trimmomatic v0.39 [49] using default parameters. Trimmed reads shorter than 50nt were discarded. Low complexity reads were discarded using *bbduk* from bbtools [50] with entropy set at 0.7 (BBMap – Bushnell B. – sourceforge.net/projects/bbmap/).

#### Read Binning

Reads were mapped to human rRNA and the human mitochondrial genome using *bbmap* from bbtools, the mapped reads were discarded. The remaining reads were binned into human genome, bacterial rRNA, and a bin that contains all microbiome reads, using *seal*, from bbtools, with default parameters. The human genome (GRCh38) and SILVA bacterial rRNA database were used as references. Binning resulted in an average of 98,120,791 (740,58,180 – median) reads mapping to the human genome and an average of 2,263,290 (848,478 – median) microbiome reads. The microbiome reads bin contains viral, bacterial, fungal, and unclassified reads.

#### Taxonomic classification of reads

Reads from the microbiome bin were subjected to taxonomic classification using KrakenUniq [51] with default parameters. The reference NCBI nt database was installed via kraken2-build script.

#### Virome Profiling

To produce a high confidence virome profile, we developed a method that first produces *de novo* transcriptome assemblies, followed by putative virome identification using BLAST searches, and finally high confidence virome profiling based on read mapping to reference virus genomes. This workflow was implemented in a bash script. First, reads that were classified as viral by KrakenUniq were extracted using the script *krakenuniq-extract-reads*, with taxonID 10239 (superkingdom, viruses). If more than 100,000 reads were extracted, they were first normalized to a target depth of 100 using *bbnorm* from bbtools. Reads were assembled using the metaSPAdes assembler. Resulting contigs were filtered for length, using *reformat* from bbtools, and only contigs that were at least 300bp were retained. Nucleotide BLAST (blastn) searches were performed on the resulting contigs, against the NCBI nt database with -max_target_seqs and -max_hsps set to 1. From the blast results, a list of subjects was compiled and their genome sequences (fasta) were extracted from the nt blast database using the blastdbcmd from BLAST. Each of those genome sequences were used as a reference and all the virome reads were mapped using bowtie2. Genome coverage and average read depth statistics were extracted from this mapping using samtools [52]. The high confidence virome profile was constructed using these coverage statistics.

#### SARS-CoV-2 genomes

Full sets of reads (before binning into human, bacterial rRNA and microbiome) were utilized to produce the SARS-CoV-2 genome sequences. They were mapped to the SARS-CoV-2 reference genome NC_045512 using bowtie2 with default parameters. Consensus sequence was called using samtools, and lineages were determined using the Pangolin web application (https://pangolin.cog-uk.io/). Consensus genome sequences were submitted to GISAID.

#### Host response to SARS-CoV-2 infection

The reads identified as originating from human transcripts were mapped to the human genome (hg19) using HISAT2 [53]. The read counts for genomic features were quantified using HTSeq [54]. The feature counts of all the samples were combined into a single matrix using a custom R script. SARS-CoV-2 positive samples were partitioned into tertiles based on CT values. Differential expression analysis was performed by comparing tertile groups using the DESeq2 package [55]. Genes with a significant log2 fold change with an adjusted p-value <0.05 were treated as differentially expressed. The lists of differentially expressed genes for each group were analyzed for enrichment of Reactome Human Pathways using Enrichr [56], and were deemed significant when FDR < 0.05.

#### Spearman Rank Correlation

Spearman’s rank correlation coefficient was calculated for gene expression counts and CT value (viral load) using the cor.test() function in R. The Benjamini-Hochberg method was used for multiple comparisons and p-value adjustments.

## Abbreviations

CT: Cycle threshold
COVID-19: Coronavirus disease 2019
SARS-CoV-2: Severe acute respiratory syndrome coronavirus-2
IQR: Interquartile range
LRT: Lower respiratory tract
RT-qPCR: Quantitative reverse transcription PCR
rRNA: Ribosomal ribonucleic acid

## Author’s contributions

SVR, JHT, EP, SM and SRD contributed to the study design. KSK, MHS, BCW, VG, MHF, and JHT contributed to the sample collection. MHS, BAS, HMB and SRD contributed to the sample processing. SBP and SVR contributed to microbiome profiling and data analysis. SBP generated SARS-CoV-2 genome sequences. SVR and MHS contributed to the statistical analysis. JHT and SRD obtained the research funding supporting this study. SVR and SRD wrote the initial version of the manuscript and all authors reviewed and approved the final version.

## Notice of Prior Presentation

None

